# Multiplexed Brain and Visceral Two-Photon Imaging Using a Simulation-Guided Ultrafast Three-Color Fiber Laser

**DOI:** 10.1101/2025.06.19.660526

**Authors:** Marvin Edelmann, Andreu Matamoros-Angles, Mohsin Shafiq, Mikhail Pergament, Franz X. Kärtner, Markus Glatzel

## Abstract

Multicolor two-photon microscopy is an essential tool in modern life sciences, enabling simultaneous, high-resolution imaging of multiple cellular structures and dynamic processes within complex biomedical systems. Realizing its full potential demands light sources that combine multiplexed spectral flexibility, high pulse quality, and practical implementation for efficient excitation of diverse cellular targets. Here, we present a novel ultrafast fiber laser platform that enables efficient three-color multiplexed two-photon imaging through numerically optimized nonlinear spectral shaping in a photonic crystal fiber (PCF). The system is driven by a nonlinear Yb-doped fiber amplifier with tailored dispersion and gain characteristics to generate clean sub-50 fs pulses at 1030 nm with over 40 nJ pulse energy. Subsequent, simulation-guided PCF-based spectral broadening enables controlled formation of three distinct high-energy bands centered at 940 nm, 1,080 nm, and 1,175 nm, overlapping with key fluorescent probes and biomolecular markers. The resulting pulses, isolated with high spectral and time-domain pulse quality, provide sub-115 fs duration and 2.5 - 6 nJ energy per channel. Multiplexed imaging is validated in labeled mouse brain, kidney, and liver tissue slices using spectrally independent multi-fluorophore targeting to visualize e.g., astrocytes, neuronal structures, and nuclei in triple-stained mouse hippocampus. The demonstrated fiber-optic laser platform provides a practical alternative to conventional single-color sources and more complex multi-laser systems, supporting robust and high-resolution three-color two-photon imaging for a range of biomedical applications.

## 1. Introduction

Two-photon microscopy (2PM) enables high-resolution, three-dimensional imaging of tissue morphology, biological dynamics, and cellular interactions at substantial depths, and is widely used in fields such as neuroscience [1], cancer research [2], and developmental biology [3]. Its capabilities can be further enhanced through spectrally multiplexed excitation and emission channels, allowing simultaneous visualization of multiple biological structures or dynamic processes. This technique, known as chromatically multiplexed two-photon microscopy (CrM-2PM), has driven numerous biomedical advances in recent years. For instance, Rakhymzhan *et al*. recently visualized germinal center reactions in mouse lymph nodes *in vivo* using CrM-2PM combined with multi-fluorophore labeling and spectral unmixing [4], while Abdeladim *et al*. achieved whole-brain, subcellular-resolution imaging in Brainbow-labeled mice to enable comprehensive mapping of neuronal circuits [5]. Given these advantages, implementing high-performance CrM-2PM remains technically challenging due to the need for spectrally distinct, ultrashort, and energetic pulses to efficiently excite multiple fluorophores [6].

Three main strategies have emerged to meet these requirements: the usage of multiple ultrafast lasers at different wavelengths [7,8], multicolor parametric wavelength conversion from a single source [9,10], or the generation of broadband pulses covering multiple excitation bands [11–13]. Among these, single broadband sources typically offer superior compactness, cost-efficiency, and alignment stability, attractive for turnkey and clinical use.

Traditionally, broadband pulse generation has relied on Ti:Sapphire lasers combined with optical parametric conversion to extend the spectral coverage [14,15]. More recently, fiber-based sources, particularly Yb-doped fiber lasers (YFLs) have emerged as robust alternatives due to their enhanced compactness, thermal efficiency, and compatibility with key fluorophores such as YFP [17] and jYCaMP6 [18]. Because of their limited gain bandwidth, broadband YFL-based sources typically combine high-energy ultrashort pulse generation with nonlinear broadening in photonic crystal fibers (PCFs) [19,20]. Combined with recent advances in low-noise, stable mode-locked fiber lasers [21], this strategy shows great promise for the realization of fiber-based CrM-2PM systems. Notably, Li *et al*. recently demonstrated a compact multicolor CrM-2PM platform using Cherenkov radiation and Raman gain in a PCF [22]. Similarly, Hsiao et al. reported a multicolor CrM-2PM source based on self-phase modulation and soliton dynamics in a PCF, generating broadband output pulses for multicolor excitation [13]. While these approaches represent significant advances in system integration and multicolor imaging capability, the broad and often uncontrolled spectral spread per wavelength channel between 150-200 nm limits the energy concentration at the desired fluorophore excitation peaks. This reduces excitation efficiency and increases the risk of out-of-band phototoxicity and spectral cross-talk between the individual color channels [23,24]. For deep-tissue 2PM applications, such broad spectral bandwidths can further accelerate temporal pulse broadening and rapid peak power degradation due to enhanced dispersion and scattering effects within the tissue [25].

To overcome these limitations, achieving greater control over the nonlinear spectral broadening process in the PCF is essential. Recent advances have focused on the precise engineering of complex nonlinear dynamics during PCF-based spectral broadening, for example through dispersive wave generation [12,26] or controlled self-phase modulation to enhance spectral shaping [27,28]. These approaches have enabled multicolor two-photon excitation and high-performance tunable single-color sources [29-32]. A key advantage of PCF-based broadening is that its nonlinear dynamics can be accurately modeled using established numerical frameworks, enabling predictive and tailored spectral design [20]. However, to our knowledge, no prior work has rigorously applied such computational optimization to engineer a fiber-based laser platform for efficient, low cross-talk three-color CrM-2PM specifically designed to target key biomedical fluorophores and biomolecular markers.

Here, we present a computationally optimized ultrafast fiber laser system for the generation of three energetic, spectrally distinct ultrashort pulses with numerically engineered energy concentration and spectral isolation, tailored for three-color biomedical CrM-2PM. By modeling both the nonlinear amplification dynamics in a pre-chirp managed Yb-doped fiber amplifier and the subsequent spectral broadening in the PCF, controlled generation of three spectrally isolated, high-energy lobes centered at 940 nm, 1,080 nm, and 1,175 nm is achieved, well-matched to key fluorophores used in modern life sciences. The resulting sub-115 fs pulses with 2.5-6 nJ of pulse energy per channel enable robust and efficient three-color excitation without the need for multiple laser sources or complex wavelength tuning mechanisms. Using this spectrally engineered three-color output, multiplexed two-photon imaging is demonstrated in triple-labeled mouse brain, kidney, and liver tissues, confirming the system’s capability to resolve complex tissue structures in diverse biological samples. This fiber-optic platform therefore combines numerically optimized three-color pulse generation with fiber-optic simplicity and versatility, offering a compact and powerful tool for advanced multicolor biomedical imaging.

## 2. Experimental Setup

As a first step, the setup of the laser source and the attached imaging system is introduced. Fig.1 (a) shows the experimentally implemented three-color fiber laser setup, developed through the numerical optimization strategy outlined in the following chapters. Initially, a homebuilt Yb-doped fiber laser (YFL), consisting of a nonlinear polarization evolution (NPE) mode-locked oscillator [21] and a single-stage preamplifier, generates a 30 MHz pulse train centered at 1,028 nm with up to 120 mW of average power. The pulse train is pre-chirped using a transmission grating pair (GP1, 1,000 lines/mm, Coherent LightSmyth 1040) to control the peak power of the pulse train before entering the subsequent pre-chirp managed fiber amplifier (PCMA) stage. This stage simultaneously amplifies and spectrally broadens the pulses via parabolic pulse evolution to achieve energetic, linearly chirped and therefore highly compressible output pulses [33-35].

**Fig 1:**
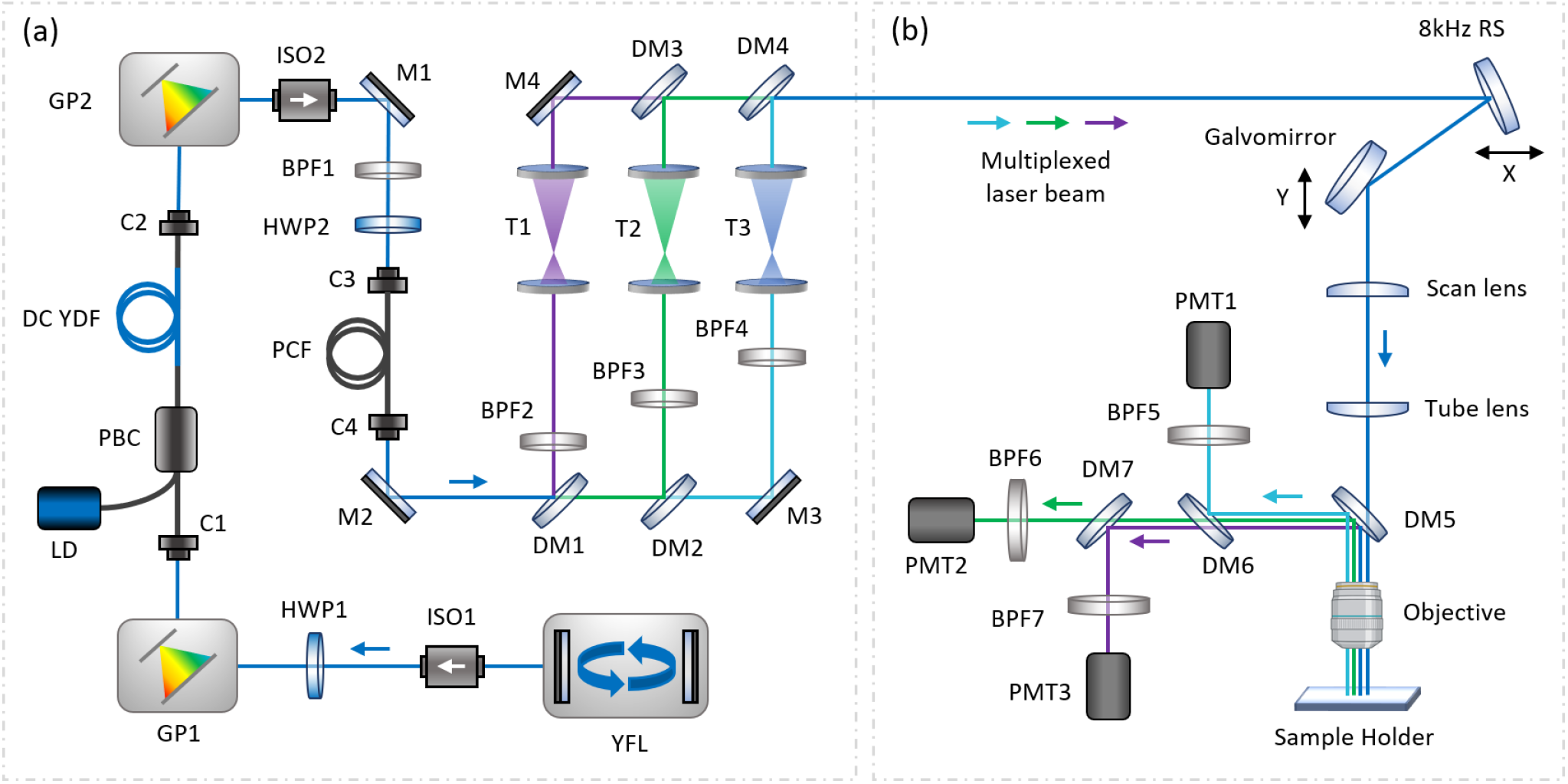
(a): Experimental setup of the three-color nonlinear Yb:fiber laser. YFL, Ytterbium fiber laser; ISO, isolator; BPF, bandpass filter; HWP, half-wave plate; GP, grating pair; C, collimator; LD, laser diode; WDM, wavelength-division multiplexer; DC YDF, double-cladding Yb-doped fiber; M, mirror; PCF, photonic crystal fiber; DM, dichroic mirror; T, telescope. (b): Experimental setup of the scanning microscope for multicolor two-photon imaging. RS, resonant scanner; PMT, photomultiplier tube.

Fiber collimators C1 and C2 (Thorlabs F240APC-980) for the coupling in and out of the PCMA. The PCMA includes 1.1 m double-cladding Yb-doped fiber (DC-YDF, Coherent PLMA-YSF-10/125). The total fiber length of the PCMA is approximately 2.4 m with 0.8 m of passive fiber (Coherent, PM1060L) at the side of C1, and 0.5 m of passive fiber at the side of C2. The YDF length is set to ensure amplifier saturation based on subsequent numerical simulations, enabling maximum efficiency and low noise performance [36]. The passive fiber lengths of the PCMA stage are kept as short as practically possible. The DC-YDF is optically pumped in the forward direction using a pump beam combiner (PBC) and an 18 W, 976 nm multimode laser diode (LD, BWT K976AA2RN).

After amplification, a second grating pair (GP2) and a subsequent 1,050±30 nm bandpass filter (BPF1) are applied for compression and additional shaping of the amplified output pulses (further details below), respectively. The pulse train is then launched into a 20 mm segment of highly nonlinear photonic crystal fiber (PCF, NKT LMA-PM-10) with up to 75% efficiency using an aspheric lens (L1, Thorlabs C240TMD-B). The PCF output beam is collimated with an achromatic lens (L2, Thorlabs AC050-008-C). Following the numerically optimized spectral broadening in the PCF, three spectral lobes are generated at center wavelengths of 950 nm, 1,080 nm, and 1,175 nm. Three-color ultrashort pulses are spectrally isolated using dichroic mirrors (DM1,2), further isolated via a set of bandpass filters (BPF2, 3, and 4), and adjusted to a ∼2.4 mm collimated beam diameter using individual telescopes (T1, 2, and 3). A second set of dichroic mirrors is then used to spatially recombine the three-color pulses at the input of the imaging system.

The 2PM setup, shown in Fig.1 (b), is based on a resonant-galvo scanning microscope (Thorlabs MPM-2PKIT) equipped with an 8 kHz resonant scanner and a 30 Hz mirror galvometer to scan the field of view (FOV) in x- and y-dimension. A water immersion objective (Olympus XLPLN25XWMP2) with 25x magnification, a numerical aperture (NA) of 1.05, and a working distance of 2 mm is used to focus the three-color multiplexed input beam onto the sample and to epi-collect the generated two-photon emission. The scanning microscope is optimized for multiplexed three-color imaging using the biomolecular markers Alexa Fluor 488 (AF488), SYTOX Orange (SO), and Alexa Fluor 647 (AF647) with emission peaks at 520 nm, 570 nm, and 670 nm, respectively. To ensure minimal spectral overlap between the channels during detection, a series of DMs is used for the initial spectral separation (DM5: 775 nm cut-on, DM6: 532 nm cut-on, DM7: 635 nm cut-on). Further spectral isolation is then ensured through the implementation of additional BPFs in front of the PMTs, aligned with the respective fluorophore emission peak (BPF5: 520±12.5 nm for AF488, BPF6: 570±16 nm for SO, BPF7: 670±12.5 nm for AF647).

The two-photon emission signal in each channel is detected using amplified GaAsP photomultiplier tubes (PMT, Thorlabs PMT2101) specified for a bandwidth of 300-720 nm with maximum sensitivity of 176 mA/W at ∼550 nm. Image acquisition and control are performed using a commercial software (Thorlabs ThorImageLS) and the system’s integrated data acquisition card (AlazarTech ATS9440).

## 3. Results and Discussion

### 3.1 Pre-chirp Managed Ytterbium Fiber Laser System

To achieve fiber-optic three-color pulse generation tailored for CrM-2PM, both the PCMA stage and the nonlinear broadening in the PCF are designed and optimized using comprehensive numerical simulations with experimental input parameters. The nonlinear propagation in the PCMA influenced by chromatic dispersion, third-order nonlinear effects from the optical Kerr-effect, and gain is numerically implemented using the nonlinear Schrödinger equation (NLSE) in the form

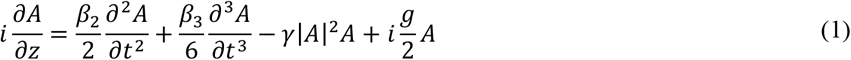

where *A*(*z, t*) denotes the complex amplitude of the electromagnetic pulse, *g* the wavelength-dependent gain factor at position *z, β*_2_ the second order group velocity dispersion (GVD) of the fiber, *β*_3_ the third-order dispersion (TOD) and *γ* = *n*_2_*ω*_0_⁄*c*_0_*A*_*eff*_ the nonlinear parameter [36]. Here, *n*_2_ is the nonlinear refractive index of the fiber material, *ω*_0_ the angular frequency of the optical carrier, *c*_0_ the speed of light in vacuum and *A*_*eff*_ the effective mode area of the fiber. Eq. (1) is solved numerically using a split-step Fourier algorithm. The gain factor *g* at each step in *z*-direction is computed by solving a two-level rate-equation for Yb-doped fibers as described in Ref. [37].

Fig.2 (a) and (b) show the measured YDF output spectrum used as simulation input and the corresponding autocorrelation (AC) trace, respectively. The AC trace is compared to the Fourier-transform limit (FTL) calculated from the measured spectrum at the input of GP1. The input pulses are positively chirped with a full width at half maximum (FWHM) of 0.7 ps assuming a Gaussian pulse-shape. The PCMA is numerically implemented according to the experimental setup with a pump power of 5.8 W. A comprehensive overview of simulation parameters is provided in Table S1 of the Supplemental Document. To generate broadband, amplified, and compressible pulses at the PCMA output, it is essential to maintain pulse evolution within the parabolic propagation regime. For fixed system parameters, this regime can be accessed by tuning the input pulse energy *E*_0_ and group-delay dispersion *GDD*_0_ i.e., the pre-chirp applied via GP1 in Fig. 1(a), numerically implemented by computing the gratings spectral phase *φ*_*GP*_(*ω*). To determine the optimal PCMA input conditions for pulse compression, *E*_0_ and *GDD*_0_ are systematically varied to maximize the Strehl-ratio, defined as the ratio of compressed to Fourier-transform limited (FTL) peak-power. Fig.2 (c) shows a simulated Strehl-ratio matrix as a function of *E*_0_ and *GDD*_0_, indicating that an average input power of 40 mW (*E*_0_ = 1.3 *nJ*) with a pre-chirp of *GDD*_0_ = −0.05 *ps*^2^ yields a maximum Strehl-ratio of up to 0.7. To enhance compression and suppress nonlinear and third-order dispersion (TOD)-induced spectral distortions, a 1,050±30 nm BPF is numerically applied using a 4^th^ order Super-Gaussian function in the frequency domain. The resulting Strehl-ratio matrix, shown in Fig.2 (c), reveals a substantial improvement with values exceeding 0.95, e.g., at ∼25 mW average power (*E*_0_ = 1.3 *nJ*), *GDD*_0_ = – 0.02 *ps*^2^ and pulse FWHM of 0.3 ps. Fig.2 (e) compares the output spectra with and without the applied BPF, while Fig.2 (f) displays the corresponding compressed output pulse, showing 46 fs FWHM and a clean temporal profile closely matching the calculated FTL pulse.

**Fig 2:**
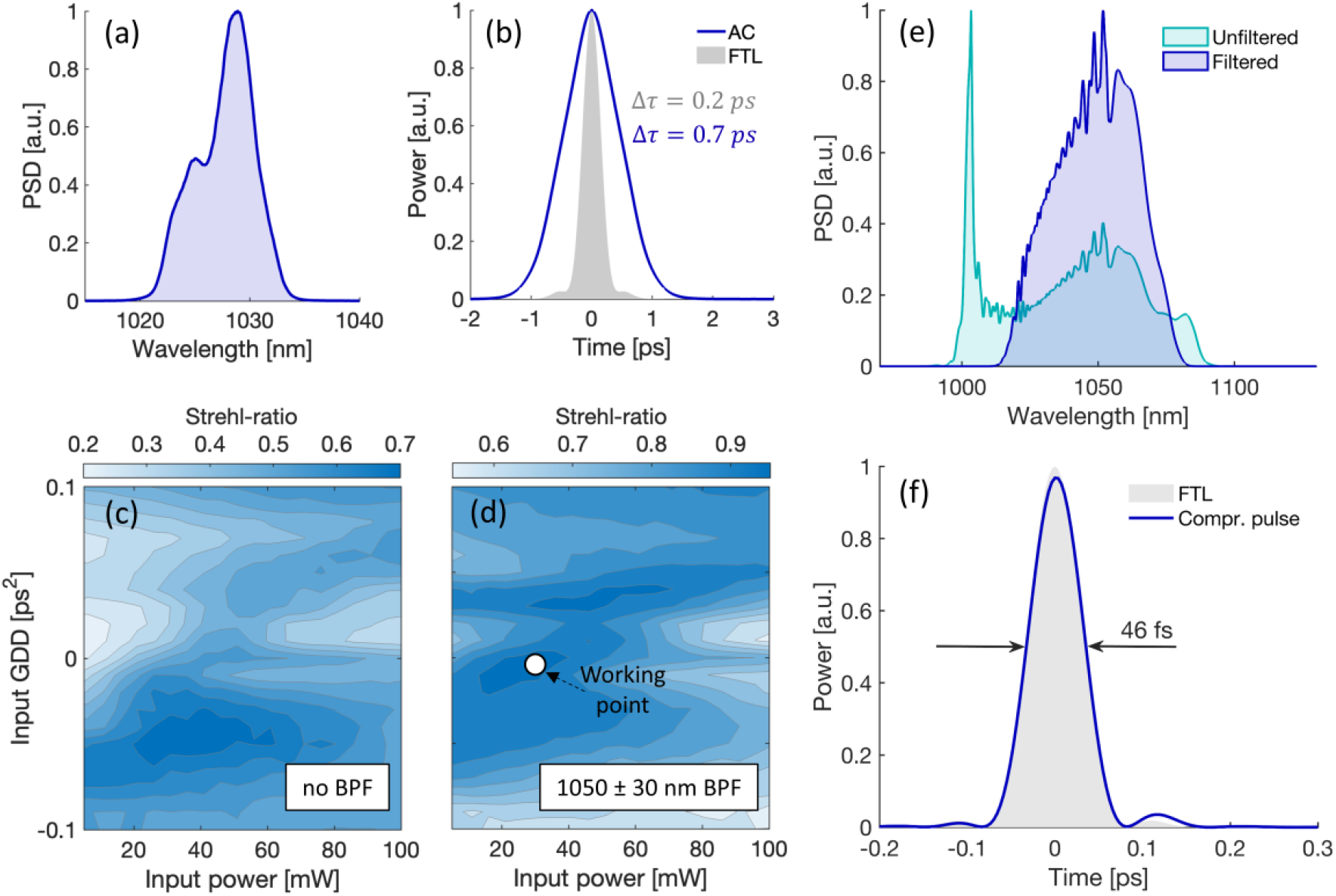
Numerical design of the PCMA-based Yb-doped fiber laser. (a): Measured YFL output spectrum used as input for the PCMA. (b): Corresponding measured AC trace compared to the calculated Fourier-transform limited (FTL) pulse. (c): Contour plot of the simulated Strehl-ratio at the PCMA output as function of the PCMA input GDD and average power without implemented BPF. (d): Strehl-ratio for identical parameter space but with implemented 1050±30 nm BPF. (e): Filtered and unfiltered output spectrum at working point marked in (d). (f): Corresponding filtered and compressed output pulse with Strehl-ratio >0.95, compared to the calculated FTL pulse.

Building on the numerical optimization results, the experimental PCMA setup shown in Fig.1 (a) is implemented to match the identified parameter space. Fig.3 (a) compares the measured output spectrum with the numerically optimized spectrum at an input average power of ∼40 mW and a pulse FWHM of 0.4 ps. The measured average output power reaches ∼1.22 W, corresponding to pulse energies exceeding 40 nJ. The strong agreement between measured and simulated spectra, both in terms of spectral shape and key features, validates the accuracy of the model. The inset of Fig.3 (a) further confirms this consistency after applying a 1,050 ± 30 nm BPF. Fig.3 (b) shows the measured autocorrelation (AC) trace of the filtered and compressed output pulse with a FWHM of 48 fs, closely matching the calculated FTL pulse width of 46 fs and the simulated pulse at the optimized Strehl-ratio working point. The successful generation of clean, sub-50 fs pulses at this energy level demonstrates the predictive power of the numerical modeling and establishes optimal conditions for the subsequent nonlinear three-color pulse generation in the PCF towards biomedical CrM-2PM application.

### 3.2 Simulation-Guided Multicolor Wavelength Conversion in a Photonic Crystal Fiber

Using the experimentally achieved ultrashort and energetic PCMA output pulses, the next step of the numerical optimization focuses on broadband nonlinear spectral broadening in a photonic crystal fiber (PCF) towards efficient multicolor-pulse generation for CrM-2PM applications. The primary target is to generate a supercontinuum with multiple, well-defined spectral peaks that align with the two-photon absorption bands of widely used fluorophores. Additionally, the resulting pulses must deliver sufficient peak power for strong two-photon signal generation, *i*.*e*., a clean pulse shape and short pulse duration (ideally sub-100 fs) with sufficient pulse energy. Regarding the targeted wavelengths, regions around 950 nm, 1,100 nm, and 1,200 nm are particularly well-suited for a broad range of imaging applications, offering both spectral compatibility with key fluorophores and sufficient separation to minimize cross-talk. For instance, 950 nm efficiently excites green-emitting fluorophores such as Alexa Fluor 488 and GFP, which is widely used e.g., in GFP-transgenic mice to visualize neural structures while preserving their function [38]. It also targets a variety of calcium indicators like Fluo-4 and jGCaMP6 for live-cell imaging of neuronal activity [39,40]. 1,100 nm aligns with yellow-to-orange fluorophores, including YFP and SYTOX Orange, commonly applied in neuronal labeling and nuclear staining, and further supports the excitation of next-generation calcium sensors such as jYCaMP [18]. The 1,200 nm band overlaps with the two-photon absorption of red and far-red dyes like Alexa Fluor 647 and ATTO 647N, ideal for deep-tissue imaging and immunolabeling.

To achieve these spectral characteristics and ensure precise control over the nonlinear dynamics, an accurate numerical model is essential. The nonlinear spectral broadening in the PCF is therefore numerically described using the generalized nonlinear Schrödinger equation (GNLSE) in the form

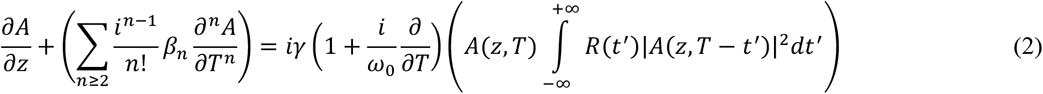

where *A*(*z, t*) again denotes the imaginary pulse envelope, *β*_*n*_ the n^th^ order dispersion coefficient, *γ* the nonlinear coefficient as used in Eq. (1), and *R*(*t*) the Raman coefficient that considers both electronic and molecular response of fused silica [20]. The GNLSE is widely used to accurately model supercontinuum generation in PCFs, considering key nonlinear effects such as self-phase modulation (SPM), self-steepening, and stimulated Raman scattering (SRS) in interaction with high-order dispersion terms, subsequently included up to the 11^th^ order.

To achieve controlled, moderate spectral broadening in the 900 -1,200 nm range, the sub-50 fs, Watt-level pulses from the PCMA are launched into a large-mode area PCF with normal (positive) dispersion, which helps to mitigate excessive nonlinear effects while supporting efficient, SPM-dominated spectral shaping of two symmetric peaks at the short and longer wavelength side. The utilized PCF (NKT, PM-LMA-10) has a zero-dispersion wavelength at ∼1300 nm (positive dispersion <1300 nm), an effective mode diameter *d*_*eff*_ = 8.5 *μm*, a NA of 0.12 at 1064 nm and a nonlinear parameter of *γ* = 3.5 *W*^−1^*m*^−1^. Table S2 of the Supplemental Document provides a detailed simulation parameter overview.

Using the measured PCMA spectrum (inset of Fig. 3(a)) with 0.64 W average power (21.8 nJ pulse energy), the PCF output spectra are simulated for pre-chirp values ranging from –0.02 ps^2^ to 0.02 ps^2^, resulting in the colormap shown in Fig.4 (a). To identify the optimal operating point for efficient three-color pulse generation, the pulse energy within 50 nm spectral bands centered at 960 nm, 1,080 nm, and 1,175 nm is analyzed as a function of the pre-chirp, as shown in Fig.4 (b). A pre-chirp of approximately 0.09 ps^2^ (corresponding to ∼75 fs input pulse duration) yields the most balanced energy distribution: ∼5.2 nJ at 960 nm, ∼3.5 nJ at 1,080 nm, and ∼2.5 nJ at 1,175 nm. The corresponding simulated PCF output spectrum at this working point is shown in Fig.4 (c), compared to the experimentally measured spectrum that is obtained using similar input parameters (0.64 W input power, ∼66 fs pulse duration). In both cases, three well-defined peaks at the target wavelengths are clearly observed with a high degree of similarity between numerical and experimental results.

**Fig 3:**
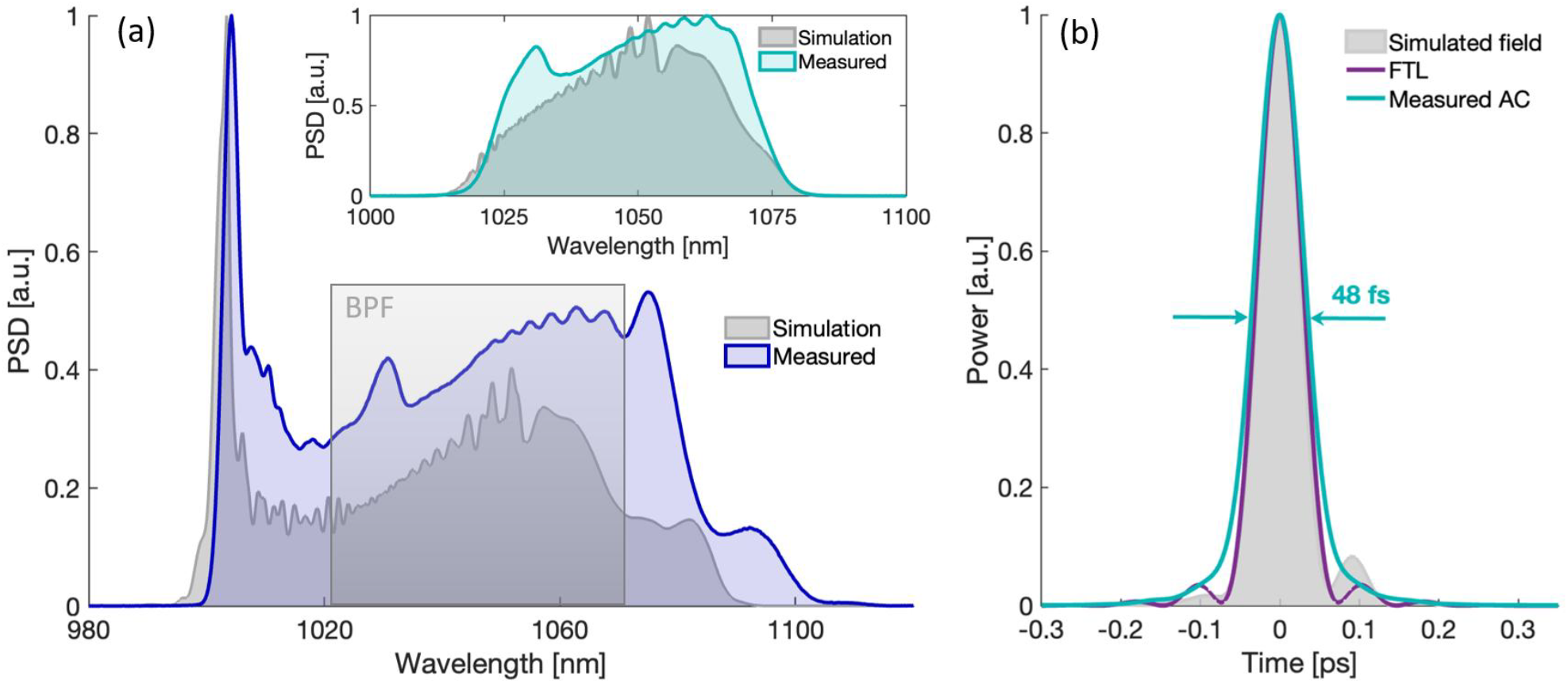
Experimental results of the PCMA-based Yb:fiber laser at the numerically determined optimal working point. (a): Measured unfiltered PCMA output spectrum in comparison to the simulated spectrum and indicated BPF application. Inset: Measured spectrum after 1050±30 nm BPF compared to the corresponding simulated spectrum. (c): Measured AC trace of the filtered and compressed pulse together with the calculated FTL pulse and the simulated output electric field.

**Fig 4:**
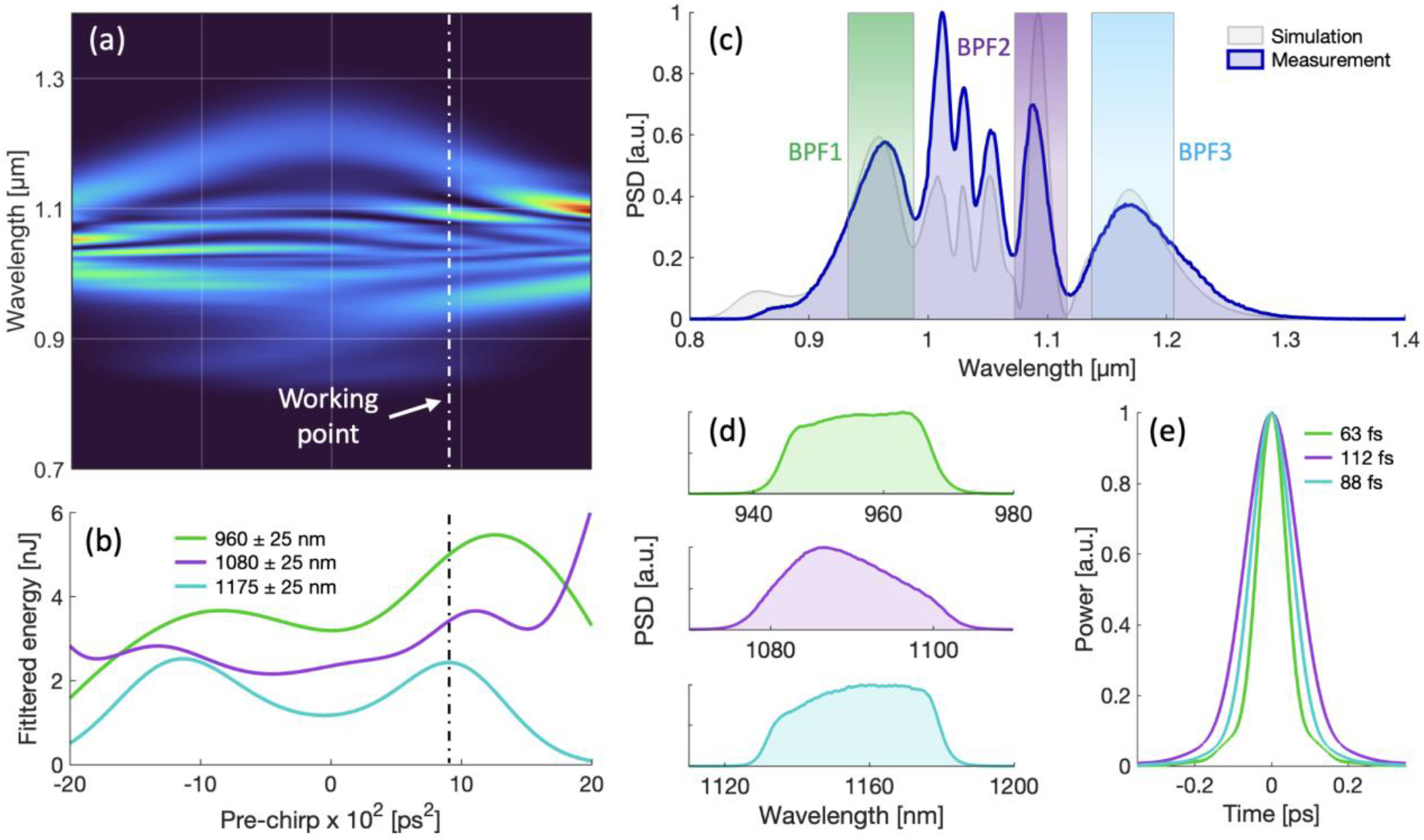
Numerical simulations and experimental results of efficient three-color generation in a PCF. (a): Simulated output spectra after 20 mm PCF for varying input pre-chirp. (b): Corresponding filtered pulse energy at 960 nm, 1080 nm and 1175 nm within a 50 nm bandwidth as function of the input pre-chirp with marked working point for efficient three-color pulse generation. (c): Simulated spectrum at the determined working point compared to the measured spectrum with indicated BPFs for color-separation. (d): Resulting measured output spectra at 960 nm (blue), 1080 nm (green) and 1175 nm (red). (e): Corresponding measured AC traces with FWHM of 63 fs (960 nm), 112 fs (1080 nm) and 88 fs (1175 nm).

Using fitting dichroic mirrors combined with the BPFs indicated in Fig.4 (c) (960±25 nm as BPF1, 1,080±12.5 nm as BPF2, and 1,175±25 nm as BPF3), the three target bandwidths are separated according to the experimental setup shown in Fig. 1 (a). At the input of the scanning microscope (i.e., considering the insertion loss of all filters and telescopes shown in Fig.1 (a)), the corresponding measured pulse energies are 3.1 nJ at 960 nm, ∼2.6 nJ at 1,080 nm and 2.8 nJ at 1,175 nm. The resulting filtered spectra are depicted in Fig.4 (d), while the corresponding AC traces in Fig. 4(e) confirm ultrashort pulse durations below 115 fs and clean temporal profiles across all three-color channels. The achieved combination of energetic, well-separated spectral bands, and sub-115 fs durations define a parameter space that is ideally suited for versatile biomedical CrM-2PM applications.

### 3.3 Three-Color Imaging of Triple-Stained Mouse Hippocampus

To demonstrate the versatility of the simulation-guided fiber laser platform for advanced biomedical imaging, three-color CrM-2PM is performed on triple-stained mouse hippocampal tissue. The staining strategy is applied to enable comprehensive, high-resolution, and simultaneous visualization of neuronal (MAP2-positive cells), glial (astrocytes, GFAP-positive cells), and nuclear components within a single tissue volume using the three-color fiber laser output. The ability to simultaneously image these components can be highly useful for a variety of neuroscientific imaging applications, *e*.*g*., for a detailed analysis of cellular architecture in healthy and diseased brain states or for *in vivo* tracing experiments of neuro-glial interactions.

Imaging is performed in the CA1 region of the dorsal hippocampus, which is involved in memory encoding, spatial navigation, and synaptic integration within hippocampal circuits [41,42]. The optical power on the sample plane is fixed at 10 mW (0.33 nJ pulse energy) per color channel to ensure sufficient signal strength with minimal photobleaching. Fig.5 (a–c) shows the acquired individual color channel images obtained from the CA1 region, recorded as the average of 3 consecutive frames in a 751×751 µm^2^ field of view (FOV), consisting of 818×818 pixels with 0.918 µm/pixel. Fig.5 (e-g) shows the same channels acquired in a posterior CA1 region using 3-frame averaging in a FOV of 532×532 µm^2^ (580×580 pixels at 0.918 µm/pixel). The corresponding channel composites for both imaged regions are shown in Fig. 5 (d) and (h), respectively. The displayed FOVs in both regions are extracted from a 940×940 µm^2^ (1024×1024 pixels at 0.918 µm/pixel) and rescaled to 1024×1024 pixels before illustration using the commercial software ImageJ. As shown, the combination of three-color laser excitation with the applied staining strategy enables high-resolution visualization of distinct pyramidal neuronal cell bodies (*stratum pyramidale, sp*) and their projections (*stratum radiatum, sr*) (cyan). In addition, individual neuronal nuclei (magenta) and surrounding astrocytes (green) are clearly resolved and spectrally distinguishable within the imaged FOVs.

**Fig 5:**
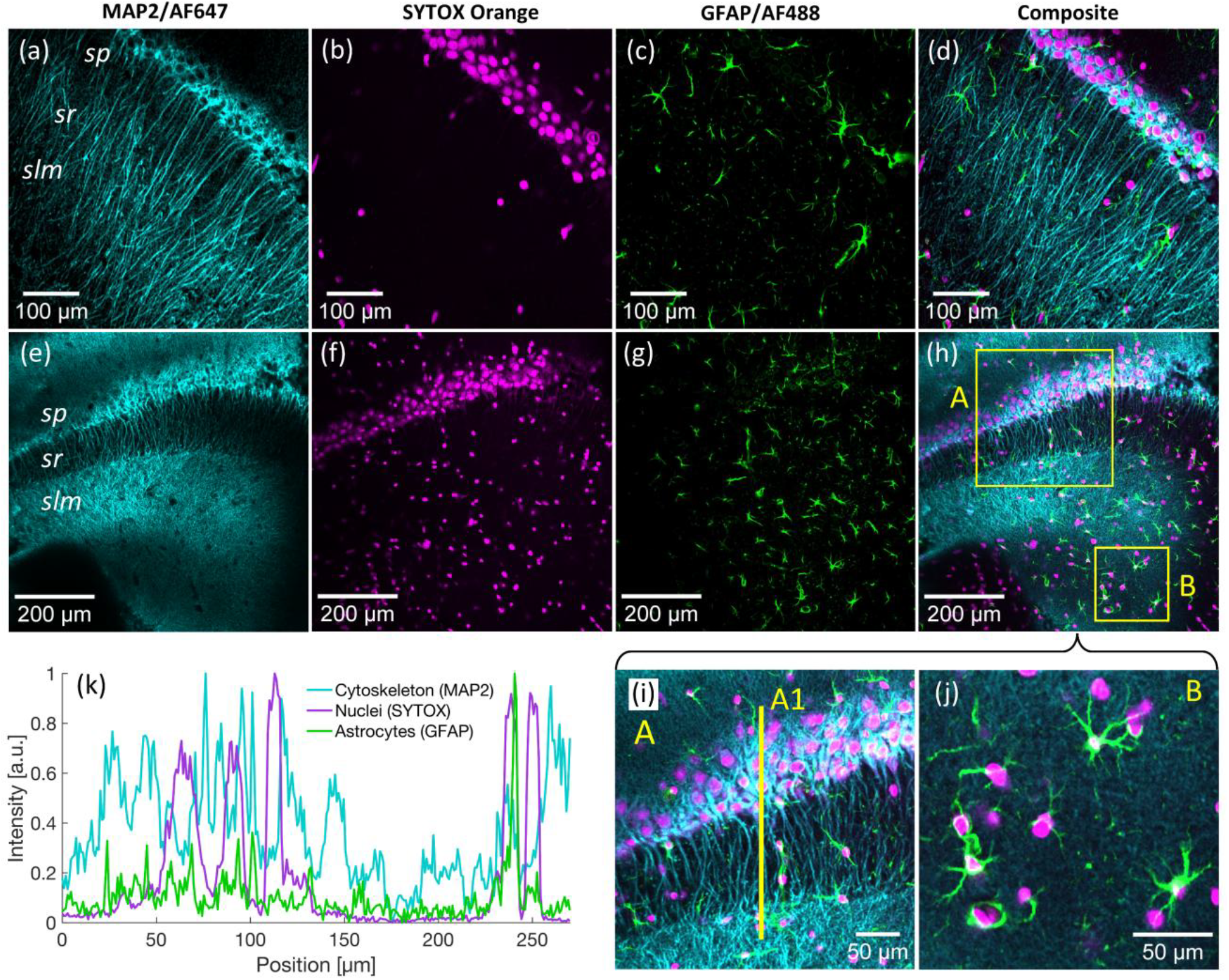
Multiplexed three-color two-photon imaging in the hippocampal region of triple-stained mouse brain. (a–d): Single-channel images in the hippocampal CA1 region showing AF647-labeled neuronal cytoskeletons (MAP2, cyan), SYTOX-labeled nuclei (magenta), and AF488-labeled astrocytes (GFAP, green), along with composite overlay. (e–h): Same channel sequence in the CA3 region with highlighted ROIs A and B in the composite channel. (i): Zoomed view of ROI A shows clear spatial separation and colocalization of astrocytes, neuronal nuclei, and MAP2-labeled dendrites; yellow line (A1) indicates the profile position. (j): Zoomed view of ROI B highlights isolated astrocytes and neuronal nuclei. (k): Intensity profile extracted along line A1, showing normalized fluorescence intensity for all three markers. *sp, sr* and *slm* states for *stratum pyramidale, stratum radiatum* and *stratum lacunosum-moleculare*, respectively.

In the composite image of the dorsal CA1 region shown in Fig.5 (h), two magnified regions of interest (ROIs A and B) are highlighted and displayed in Fig.5 (i) and Fig.5 (j), respectively. ROI A, corresponding to a 340 × 340 µm^2^ FOV (333 × 333 pixels at 0.918 µm/pixel), captures a densely populated pyramidal layer with strongly aligned neuronal soma, associated nuclei, and dendritic projections. The structural organization of astrocytes embedded between neuronal elements is clearly visible, highlighting the complementary spatial arrangement of glial and neuronal components. ROI B, representing a less densely populated region in a 187×187 µm^2^ FOV (204×204 pixels at 0.918 µm/pixel), shows isolated and clearly resolved astrocytes extending between sparsely distributed nuclei and fibers, effectively demonstrating the system’s ability to resolve fine cellular details and local interactions across all three color-channels.

Neuronal cytoskeletons and astrocytes are visualized using antibodies against Microtubule-Associated Protein 2 (MAP2) and Glial Fibrillary Acidic Protein (GFAP), conjugated to Alexa Fluor 647 and Alexa Fluor 488 with efficient two-photon excitation at 1,175 nm and 940 nm, respectively. MAP2 highlights the neuronal cytoskeleton, mainly labeling the dendrites and axons, while GFAP selectively targets the intermediate filaments of the astrocytes’ cytoskeleton. A third channel is introduced using the cell-impermeant nucleic acid stain SYTOX Orange to label the nuclei of all the cells in the tissue (including neurons, astrocytes, but also microglia and endothelial cells), excited efficiently at 1,080 nm via the central spectral lobe of the PCF-broadened output.

To quantitatively assess the three-color imaging performance of the laser and imaging setup, Fig.5 (k) further shows the intensity profile extracted along line A1, as indicated in ROI A (Fig.5 (i)). The plot reveals distinct spatial distributions of SYTOX-labeled nuclei, MAP2-labeled cytoskeletal processes, and GFAP-labeled astrocytes. Sharp peaks in the SYTOX channel reflect densely packed neuronal soma, while the broader MAP2 signal corresponds to surrounding dendritic structures. GFAP intensity peaks appear adjacent to or between neuronal layers, consistent with the expected spatial organization of astrocytes enveloping neurons. The minimal overlap between intensity peaks and high signal-to-background contrast confirm effective spectral separation, low cross-excitation, and robust fluorophore-target labeling, validating the laser systems’ capability for simultaneous CrM-2PM imaging of multiple, spatially distinct cellular structures.

### 3.4 Three-Color Imaging in Labeled Mouse Kidney and Liver Tissue

To further demonstrate the versatility and broad applicability of the three-color fiber laser system across diverse tissue types, we next applied three-color CrM-2PM to structurally labeled mouse liver and kidney tissue. The used reference samples (FluoTissue, SunJin Lab) are mounted on a prepared microchamber slide in an aqueous tissue clearing reagent (RapiClear 1.52, SunJin Lab) and stained with AF488 and SYTOX Orange to label blood vessels and nuclei, respectively. Additionally, anti-TUJ1 (anti–βIII-tubulin) antibodies with conjugated AF647 are used to label the neurons of the peripheral nervous system. Imaging is performed using the three-color fiber laser system with again ∼10 mW average optical power on the sample per channel for maximized two-photon fluorescence while avoiding photobleaching. Fig.6 shows images of the triple-stained mouse liver and kidney tissue sections, obtained in a 940×940 µm FOV (1,024×1,024 pixels at 0.918 µm/pixel) as the average of 3 consecutive frames.

**Fig 6:**
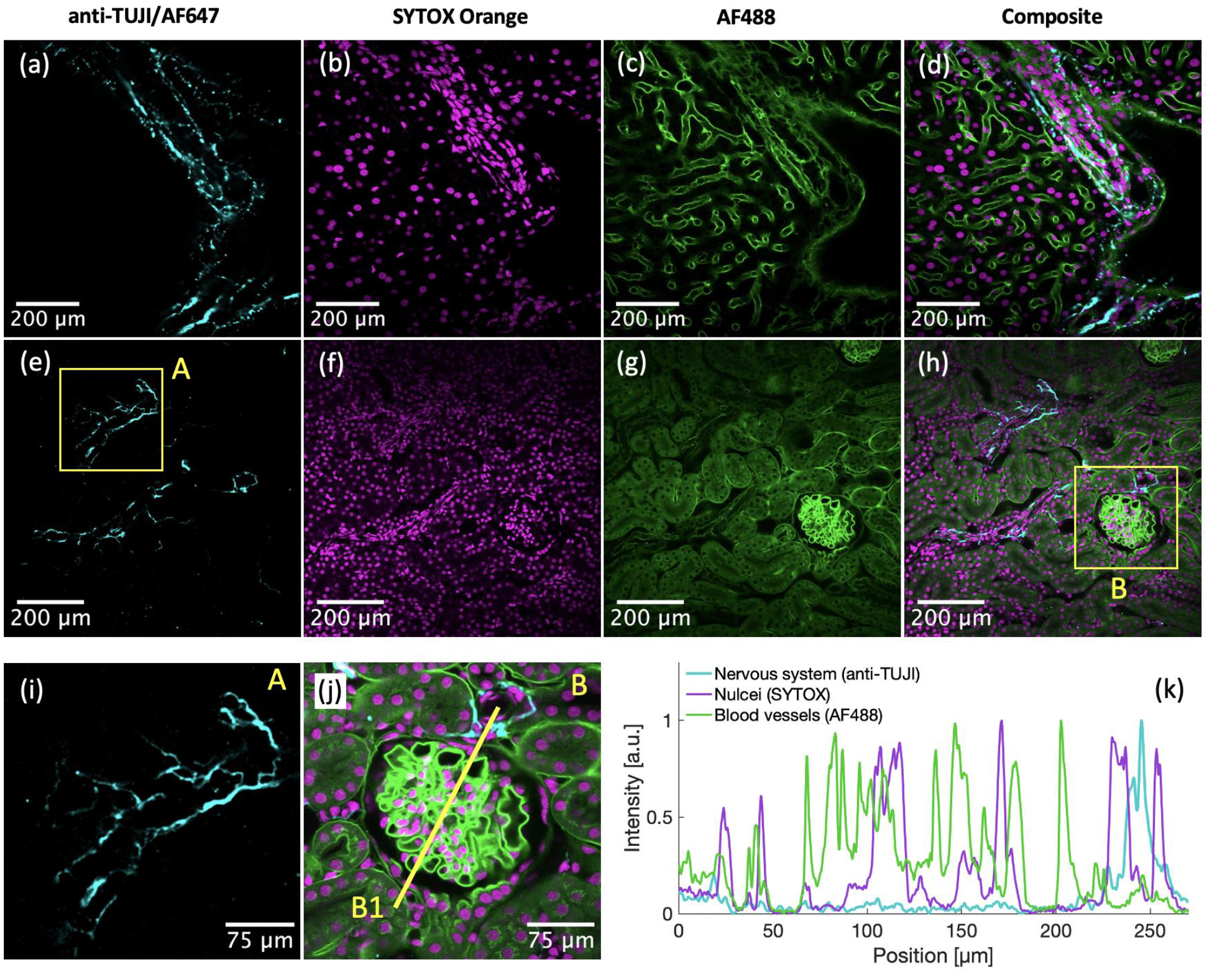
Multiplexed three-color two-photon imaging in triple-stained mouse liver and kidney tissue. (a–d): Single-channel images in mouse liver showing the AF647-labeled peripheral nervous system (anti-TUJI, cyan), SYTOX-labeled nuclei (magenta), and AF488-labeled blood vessels (green), along with composite overlay. (e–h): Same channel sequence in mouse kidney with highlighted ROIs A and B in the anti-TUJI and composite channel, respectively. (i): ROI A showing a clear visualization of individual nerve projections. (j): ROI B highlighting an isolated glomerulus with conjugated nerves and nuclei. (B1) indicates the line profile position. (k): Intensity profile along line B1, showing the normalized fluorescence intensity for all three markers.

Images obtained from the liver tissue are shown in Fig.6 (a-d): individual fluorescence channels corresponding to neuronal structures (anti-TUJ1/AF647, cyan), nuclei (SYTOX Orange, magenta), and vasculature (AF488, green) are displayed in (a–c), with the merged composite image in (d) illustrating the spatial relationships between these components. Similarly, Fig.6 (e–h) present kidney tissue images acquired using the same three-color approach, with single channels shown in (e–g) and the composite overlay in (h). Two magnified areas corresponding to the marked ROIs A and B in Fig.6 (e) and (h) are shown in Fig.6 (i) and (j), respectively. ROI A, shown in a digitally magnified 323×323 µm FOV (352×352 pixels at 0.918 µm/pixel), demonstrates the resolution of fine nerve fiber networks, highlighting the ability of the imaging system to resolve intricate neuronal structures with high spatial detail. ROI B, shown in a digitally zoomed-in 330.5×330.5 µm FOV (360×360 pixels at 0.918 µm/pixel), provides a closer view of an isolated glomerulus, where the spatial relationship between vasculature, nuclei, and nerve fibers is clearly visualized to demonstrate the system’s capability for high-resolution, multi-color imaging in dense and complex tissue environments. To further quantify the spatial distribution of the different tissue components, a fluorescence intensity profile along the marked yellow line B1 in Fig.6 (j), representing ROI B, is depicted in Fig.6 (k). Similar to the performance of the previous neuro-glial imaging experiment, distinct and well-separated peaks clearly verify the imaging system’s sensitivity and contrast without notable channel cross-talk, enabling clear differentiation between the anatomical features *in situ*.

## 4. Conclusion

In conclusion, this work presents a compact, simulation-guided ultrafast fiber laser system that, enables efficient three-color two-photon excitation through tailored nonlinear spectral broadening in a photonic crystal fiber. Numerical engineering of both the nonlinear amplification and subsequent spectral shaping enables the generation of three spectrally distinct ultrashort pulses centered at 940 nm, 1,080 nm, and 1,175 nm, with pulse energies (>2.5 nJ) and durations (<115 fs) well-suited for high-contrast, multichannel two-photon imaging. The resulting turnkey laser setup enables simultaneous, spectrally separated excitation of widely applied fluorophores without the need for multiple laser sources, additional parametric wavelength conversion or wavelength-tuning mechanisms. Experimental validation through multiplexed imaging of triple-stained mouse brain, kidney, and liver tissues confirms the system’s capability to resolve diverse cellular and subcellular features in densely structured biological systems with high spatial resolution and minimal spectral cross-talk. These results establish a robust and versatile fiber-based imaging platform with broad applicability across neuroscience, cancer research, and systems biology, representing a significant step towards the applicability of fiber-optic multicolor laser systems in biomedical imaging applications.

## A. Methods Animals

The mouse model is an adult wild-type C57BL/6J male mouse (∼6 months old, P180–P210) with ubiquitous EGFP expression. No experiments with living animals were performed. The animal breeding and euthanasia to obtain the tissue were approved by the ethical research committees of respective national/local authorities: *Freie und Hansestadt Hamburg, Behörde für Gesundheit und Verbraucherschutz*, Hamburg, Germany (ORG1108). Sampling was performed following the protocols and guidelines of the Ethical Committee and the directives of the European Union Council 86/609 and 2010/63.

### Sample Preparation

The brain of the adult mouse model was quickly extracted from the skull and fixed with 4% paraformaldehyde for 24h in the fridge. The following day, the tissue was washed with 1 M phosphate buffer (PBS) three times. Subsequently, the brain was sliced coronally (∼2 mm thick) with a scalpel, and then, under the microscope the hippocampus area was separated and placed in a compartment of a glass-bottom µ-slide microscopy chamber (Ibidi, #80427-90). Then, the tissues were impermeabilized with 0.2% Triton (in PBS) for 1h. Afterward, after 2 washes with PBS, the primary antibodies were applied in PBS for 4 days at 4°C (shaking smoothly): MAP-2 (1:300, #188002, Synaptic Systems) and (1:300, #173011, Synaptic Systems). Then, after two long PBS washes, the secondary antibodies were applied in PBS (+ 10 % Goat Serum) for 3 days: Alexa anti-Rabbit 488 (1:300, #A21202 Invitrogen) and Alexa anti-Mouse 647 (1:300, #A21443 Invitrogen). The nuclei dye SYTOX Orange (1:10,000, #S11368, Thermo Fisher) was applied together with the secondary antibodies. Finally, 3 washes with PBs were applied, and the sample was mounted using Fluoromount (Invitrogen, #00-4958-02). Imaging was performed through the ∼0.5 mm thin undersurface of the slide, to enable the insertion of a purified water droplet between the sample and the water immersion objective.

## B. Funding

This work has been supported by Deutsches Elektronen-Synchrotron DESY, a member of the Helmholtz Association (HGF), POF IV DMC and the Cluster of Excellence ‘Advanced Imaging of Matter’ of the Deutsche Forschungsgemeinschaft (DFG) – EXC 2056 – project ID 390715994. M.S.’s contribution was supported by the PIER Hamburg/Boston seed grant (PHM-2019-03) and PIER seed grant (PIF 2020-10) awarded by the University of Hamburg, Germany; and the Forschungsförderungsfonds der Medizinischen Fakultät grant (NWF-20/10) awarded by the University Medical Center Hamburg-Eppendorf (UKE), Germany. Additionally, M.G. would like to express their gratitude for the financial support received from the Joachim Herz Stiftung in Hamburg, Germany.

## C. Acknowledgements

The authors would like to thank Edda Thies from the UKE Institute of Neuropathology for assistance in sample preparation and staining.

## D. Competing Interest

The authors declare no competing interests.

## E. Data availability

Data underlying the results presented in this paper are not publicly available at this time but may be obtained from the corresponding author upon reasonable request.

## F. Author Contributions

M.E., A.M.-A. and M.S. conceived and conceptualized the experiments. M.E. conceived and constructed the laser system and performed the laser characterization and imaging experiments. A.M.-A. and M.S. prepared the triple-stained mouse brain samples. M.G., F.X.K and M.P. supervised the project. M.E. wrote the manuscript with contributions from all coauthors.

## Notes

### Competing Interest Statement

The authors have declared no competing interest.

## References

1. C. Grienberger, A. Giovannucci, W. Zeiger, et al., “Two-photon calcium imaging of neuronal activity,” Nat. Rev. Methods Primers 2, 67 (2022).

2. S. W. Perry, R. M. Burke, and E. B. Brown, “Two-photon and second harmonic microscopy in clinical and translational cancer research,” Ann. Biomed. Eng. 40, 277–291 (2012).

3. Qiang Huang, M. A. Cohen, F. C. Alsina, et al., “Intravital imaging of mouse embryos.,” Science 368,181 186 (2020).

4. A. Rakhymzhan, R. Leben, H. Zimmermann, et al., “Synergistic strategy for multicolor two-photon microscopy: application to the analysis of germinal center reactions in vivo,” Sci. Rep. 7, 7101 (2017).

5. L. Abdeladim, K. S. Matho, S. Clavreul, et al., “Multicolor multiscale brain imaging with chromatic multiphoton serial microscopy,” Nat. Commun. 10, 1662 (2019).

6. F. Helmchen, W. Denk, “Deep tissue two-photon microscopy,” Nat Methods 2, 932–940 (2005).

7. J. Herz, V. Siffrin, A. E. Hauser, A. U. Brandt, et al., “Expanding two-photon intravital microscopy to the infrared by means of optical parametric oscillator,” Biophys. J. 98, 715–723 (2010).

8. R. Galli, T. Siciliano, D. Aust, S. Korn, et al., “Label-free multiphoton microscopy enables histopathological assessment of colorectal liver metastases and supports automated classification of neoplastic tissue,” Sci. Rep. 13, 4274 (2023).

9. K. Wang, T. Liu, J. Wu, N. G. Horton, et al., “Three-color femtosecond source for simultaneous excitation of three fluorescent proteins in two-photon fluorescence microscopy,” Biomed. Opt. Express 3, 1972–1977 (2012).

10. K. Guesmi, L. Abdeladim, S. Tozer, P. Mahou, et al. “Dual-color deep-tissue three-photon microscopy with a multiband infrared laser,” Light Sci. Appl. 7, 12 (2018).

11. R. S. Pillai, C. Boudoux, G. Labroille, N. Olivier, et al., “Multiplexed two-photon microscopy of dynamic biological samples with shaped broadband pulses,” Opt. Express 17, 12741–12752 (2009).

12. L. Chou, S. Wu, H. Hung, W. Lin, et al., “Compact multicolor two-photon fluorescence microscopy enabled by tailorable continuum generation from self-phase modulation and dispersive wave generation,” Opt. Express 30, 40315–40327 (2022).

13. Y. Hsiao, Y. Huang, B. Jyoti Borah, S. Chen, et al., “Single-laser-based simultaneous four-wavelength excitation source for femtosecond two-photon fluorescence microscopy,” Biomed. Opt. Express 12, 4661–4679 (2021).

14. M. H. Brenner, D. Cai, J. A. Swanson, and J. P. Ogilvie, “Two-photon imaging of multiple fluorescent proteins by phase-shaping and linear unmixing with a single broadband laser,” Opt. Express 21, 17256–17264 (2013).

15. P. Mahou, M. Zimmerley, K. Loulier, K. S. Matho et al., “Multicolor two-photon tissue imaging by wavelength mixing,” Nat. Methods 9, 815–818 (2012).

16. Wei Shi, Qiang Fang, Xiushan Zhu, R. A. Norwood, and N. Peyghambarian, “Fiber lasers and their applications [Invited],” Appl. Opt. 53, 6554–6568 (2014).

17. E. P. Perillo, J. E. McCracken, D. C. Fernée, J. R. Goldak, et al., “Deep in vivo two-photon microscopy with a low cost custom built mode-locked 1060 nm fiber laser,” Biomed. Opt. Express 7, 324–334 (2016)

18. M. A. Mohr, D. Bushey, A. Aggarwal, J. S. Marvin, et al., “jYCaMP: an optimized calcium indicator for two-photon imaging at fiber laser wavelengths,” Nat. Methods 17, 694–697 (2020).

19. P. Russell, “Photonic Crystal Fibers,” Science 299,358-362 (2003).

20. J. M. Dudley, G. Genty, and S. Coen. “Supercontinuum generation in photonic crystal fiber.” Rev. Mod. Phys. 78, 1135–1184 (2006).

21. J. Kim and Y. Song, “Ultralow-noise mode-locked fiber lasers and frequency combs: principles, status, and applications,” Adv. Opt. Photon. 8, 465–540 (2016).

22. K. Li, L. L. H. Huang, J. Liang, and M. Chan, “Simple approach to three-color two-photon microscopy by a fiberoptic wavelength convertor,” Biomed. Opt. Express 7, 4803–4815 (2016).

23. P. Luu, S.E. Fraser, and F. Schneider, “More than double the fun with two-photon excitation microscopy,” Commun. Biol. 7, 364 (2024).

24. I. Pastirk, J. M. Dela Cruz, K. A. Walowicz, V. V. Lozovoy, and M. Dantus, “Selective two-photon microscopy with shaped femtosecond pulses,” Opt. Express 11, 1695–1701 (2003).

25. X. Liang, W. Hu, and L. Fu, “Pulse compression in two-photon excitation fluorescence microscopy,” Opt. Express 18, 14893–14904 (2010).

26. M. Edelmann, A. Matamoros-Angles, M. Shafiq, M. Pergament, et al., “Deep-Tissue Two-Photon Brain Imaging Enabled by a Tunable Fiber-Optic Dispersive Wave Generator.” bioRxiv, 2025-03 (2025).

27. W. Liu, C. Li, Z. Zhang, F. X. Kärtner, and G. Chang, “Self-phase modulation enabled, wavelength-tunable ultrafast fiber laser sources: an energy scalable approach,” Opt. Express 24, 15328–15340 (2016).

28. G. Zhou, Q. Cao, F. X. Kärtner, and G. Chang, “Energy scalable, offset-free ultrafast mid-infrared source harnessing self-phase-modulation-enabled spectral selection,” Opt. Lett. 43, 2953–2956 (2018).

29. W. Liu, S. Chia, H. Chung, R. Greinert, et al., “Energetic ultrafast fiber laser sources tunable in 1030–1215 nm for deep tissue multi-photon microscopy,” Opt. Express 25, 6822–6831 (2017).

30. H. Chung, R. Greinert, F. X. Kärtner, and G. Chang, “Multimodal imaging platform for optical virtual skin biopsy enabled by a fiber-based two-color ultrafast laser source,” Biomed. Opt. Express 10, 514–525 (2019).

31. J. R. Unruh, E. Shane Price, R. Gagliano Molla, L. Stehno-Bittel, et al., “Two-photon microscopy with wavelength switchable fiber laser excitation,” Opt. Express 14, 9825–9831 (2006).

32. Y. Liu, H. Tu, W. A. Benalcazar, E. J. Chaney and S. A. Boppart, “Multimodal Nonlinear Microscopy by Shaping a Fiber Supercontinuum From 900 to 1160 nm,” IEEE J. Sel. Top. Quantum Electron. 18, 1209–1214 (2012).

33. V. I. Kruglov, A. C. Peacock, J. M. Dudley, and J. D. Harvey, “Self-similar propagation of high-power parabolic pulses in optical fiber amplifiers,” Opt. Lett. 25, 1753–1755 (2000).

34. M. E. Fermann, V. I. Kruglov, B. C. Thomsen, J. M. Dudley, and J. D. Harvey, “Self-Similar Propagation and Amplification of Parabolic Pulses in Optical Fibers,” Phys. Rev. Lett. 84, 6010–6013 (2000).

35. Y. Deng, C.-Y. Chien, B. G. Fidric, and J. D. Kafka, “Generation of sub-50 fs pulses from a high-power Yb-doped fiber amplifier,” Opt. Lett. 34, 3469–3471 (2009).

36. G. P. Agrawal, Nonlinear Fiber Optics (Academic Press, 2007).

37. R. Paschotta, J. Nilsson, A. C. Tropper, and D. C. Hanna, “Ytterbium-doped fiber amplifiers,” IEEE J. Quantum Electron. 33, 1049–1056 (1997).

38. W. Wei, J. Elstrott, and M. B. Feller. “Two-photon targeted recording of GFP-expressing neurons for light responses and live-cell imaging in the mouse retina.” Nat. Protocols 5, 1347–1352 (2010).

39. C. Grienberger and A. Konnerth. “Imaging calcium in neurons.” Neuron 73 (2012): 862–885.

40. C. Grienberger, A. Giovannucci, W. Zeiger, and C. Portera-Cailliau, “Two-photon calcium imaging of neuronal activity,” Nat. Rev. Methods Primers 2, 67 (2022).

41. C. Dong, A.D. Madar, and M. E. J. Sheffield, “Distinct place cell dynamics in CA1 and CA3 encode experience in new environments,” Nat. Commun. 12, 2977 (2021).

42. P. E. Gilbert, and A. M. Brushfield. “The role of the CA3 hippocampal subregion in spatial memory: a process oriented behavioral assessment.” Prog. Neuropsychopharmacol. Biol. Psychiatry 33, 774–781 (2009).

