## Supplementary Table S1, Supplementary Table S2 for "Multiplexed Brain and Visceral Two-Photon Imaging Using a Simulation-Guided Ultrafast Three-Color Fiber Laser"

### Table S1. Numerical Parameters for PCMA Simulations

| Parameter | Symbol | Value |
|---|---|---|
| **Input Pulse (Measured from YDF):** | | |
| Pulse energy | $E_0$ | Variable as shown in Fig. 2 (a) and (b) |
| Repetition rate | $f_{rep}$ | 30 MHz |
| Group-delay dispersion (Pre-chirp) | $GDD_0$ | Variable as shown in Fig. 2 (a) and (b) |
| Center wavelength | $\lambda_c$ | 1050 nm |
| Spectral bandwidth (FWHM) | $\Delta\lambda$ | ~30 nm |
| **PCMA Parameters:** | | |
| Passive fiber length at C1 | $L_{p1}$ | 0.8 m |
| Passive fiber length at C2 | $L_{p2}$ | 0.5 m |
| Gain fiber length | $L_g$ | 1.1 m |
| Mode-field diameter | MFD | 10.5 µm @ 1060 nm (both gain and passive fiber) |
| Nonlinear refractive index | $n_2$ | $2.6 * 10^{-20}$ $m^2/W$ |
| Group-velocity dispersion | $\beta_2$ | 23 $fs^2/mm$ |
| Third-order dispersion | $\beta_3$ | 46 $fs^2/mm$ |
| Doping concentration | - | $1.1 * 10^8$ $\mu m^{-3}$ |
| Cross-sections | - | R. Paschotta, et al, IEEE Journal of quantum electronics, Vol.33, p 1049 (1997). |
| Pump loss | - | Y. Hua, et al., Opt. Express 26, 6427-6438 (2018). |
| **Grating compressor** | | |
| Angle of incidence | - | 31.3° |
| Line density | - | 1000 lines/mm |
| Grating distance | $d_g$ | Variable to optimize Strehl-ratio |
| **Bandpass filter** | | |
| Center wavelength | $\lambda_{c,BPF}$ | 1048 nm |
| Spectral bandwidth (FWHM) | $\Delta\lambda_{BPF}$ | 60 nm |
| Super-Gaussian order | - | 4 |

**Table S2. Numerical Parameters for PCF Broadening Simulations**

| Parameter | Symbol | Value |
|---|---|---|
| **Input Pulse (Measured from filtered PCMA output):** | | |
| Pulse energy | $E_0$ | 21.8 nJ |
| Repetition rate | $f_{rep}$ | 30 MHz |
| Temporal FWHM | $\Delta t$ | 62 fs |
| Group-delay dispersion (pre-chirp) | $GDD_0$ | $9 * 10^2$ $fs^2$ |
| Third order dispersion | $TOD_0$ | $-0.6 * 10^4$ $fs^3$ |
| Center wavelength | $\lambda_c$ | 1048 nm |
| Spectral bandwidth (FWHM) | $\Delta\lambda$ | ~50 nm |
| **PCF Parameters** | | |
| Mode field diameter | $d_{eff}$ | 8.5 µm @ $\lambda_c$ |
| Fiber length | $L_{PCF}$ | 20 mm |
| Nonlinear refractive index | $n_2$ | $33.8 * 10^{-20}$ $m^2/W$ |
| Dispersion profile | - | Imported from datasheet (NKT, PM-LMA-10) |
| Numerical aperture | NA | 0.12 @ 1064 nm |
| Zero dispersion wavelength | $\lambda_{zdw}$ | 1300 nm |
